# Three recent sex chromosome-to-autosome fusions in a *Drosophila virilis* strain with high satellite content

**DOI:** 10.1101/2021.06.14.448339

**Authors:** Jullien M. Flynn, Kevin B. Hu, Andrew G. Clark

## Abstract

The karyotype, or number and arrangement of chromosomes, has varying levels of stability across both evolution and disease. Karyotype changes often originate from DNA breaks near the centromeres of chromosomes, which generally contain long arrays of tandem repeats or satellite DNA. *Drosophila virilis* possesses among the highest relative satellite abundances of studied species, with almost half its genome composed of three related 7bp satellites. We discovered a strain of *D. virilis* that we infer recently underwent three independent chromosome fusion events involving the X and Y chromosomes, in addition to one subsequent fission event. Here we isolate, characterize and propose a timeline for the chromosome fusions in this strain which we believe demonstrates a remarkable karyotype instability. We discovered that one of the substrains with an X-autosome fusion has a X-to-Y chromosome nondisjunction rate 20x higher than the *D. virilis* reference strain (21% vs. 1%). Finally, we found an overall higher rate of DNA breakage in the substrain with higher satellite DNA compared to a genetically similar substrain with less satellite DNA. This suggests satellite DNA abundance may play a role in the risk of genome instability. Overall, we introduce a novel system consisting of a single strain with four different karyotypes, which we believe will be useful for future studies of genome instability, centromere function, and sex chromosome evolution.

## INTRODUCTION

Genome instability is characterized by large-scale mutations or rearrangements that result from repair of DNA breaks, and is a fundamental driver of karyotype evolution, chromosomal disorders, and cancer rearrangements (Black and Giunta 2018; Mayrose and Lysak 2021). The initiating event is spontaneous DNA damage such as double-stranded DNA breaks, one of the most dire events to occur in a cell (Featherstone and Jackson 1999). Even if DNA breaks are repaired, they often result in large-scale genome rearrangements. Robertsonian translocations, one of the most common rearrangements in medical genetics and evolution, occur when there are breaks near the centromere of acrocentric chromosomes and when they are repaired they are fused to each other (Mayrose and Lysak 2021). Robertsonian translocations are associated with multiple miscarriages in humans and aneuploidy disorders like Patau and Down syndromes (Braekeleer and Dao 1990), with increased rates of aneuploidy driven by increased nondisjunction of Robertsonian or fused chromosomes (Schulz *et al.* 2006). Spontaneous Robertsonian translocation events, or chromosome fusions, are rarely detected in experimentally-tractable systems. Several consecutive steps must occur in order to detect a spontaneous fusion: 1) DNA breakage near the centromere; 2) repair and fusion with another acrocentric chromosome; 3) resolution of a single functional centromere; and 4) retention and increase in frequency of the fusion in the line. Natural genetic variation and environmental conditions that can affect the rate or probability of any of the steps are not understood.

Satellite DNA consists of long arrays of tandemly repeated sequences, and is often located near centromeres in heterochromatin (reviewed in Thakur *et al.* 2021). Satellite DNA varies greatly in sequence and abundance within and between species (Subirana *et al.* 2015; Wei *et al.* 2018; Cechova *et al.* 2019). Although satellite DNA differences between some species have been linked to reproductive incompatibilities (Ferree and Barbash 2009; Jagannathan and Yamashita 2021), the biological implications of intraspecies abundance variation has not been explored. Satellite DNA can vary in abundance by several megabases among individuals of the same species, including in flies and humans (Miga *et al.* 2014; Wei *et al.* 2014; Flynn *et al.* 2020). Satellite DNA appears to be constrained by maximum limits, with no species studied so far having more than about half of their genome made up of satellite DNA (Gall and Atherton 1974a; Fry and Salser 1977; Petitpierre *et al.* 1995). In past modeling efforts, satellite DNA arrays have been proposed to be weakly deleterious until they reach a maximum length beyond which they are not tolerated by selection(Charlesworth *et al.* 1986). Slow DNA replication or development time have been suggested as mechanisms to enforce strong negative selection against long satellite arrays, however empirical evidence for this has been limited (but see Bilinski *et al.* 2018). Most karyotype rearrangements detected in both humans and other organisms originated from breaks near the centromere, especially in or near satellite DNA (Black and Giunta 2018; Balzano *et al.* 2020). This may be due to intrinsic instability of satellite DNA arrays caused by replication stress of polymerases progressing through highly repetitive sequences, or the formation of unstable DNA topology (Barra and Fachinetti 2018).

*Drosophila virilis* is an excellent model for studying satellite DNA variation and karyotype stability. *D. virilis* has the highest relative abundance of simple satellite DNA (defined as satellites with unit length <=20 bp) compared to any other studied species. Three 7 bp satellites, AAACTAC, AAACTAT, and AAATTAC take up over 40% of the genome in *D. virilis,* and they form arrays tens of megabases long in the pericentromeric region (Gall *et al.* 1971; Gall and Atherton 1974b; Flynn *et al.* 2020). The *D. virilis* reference strain has retained the ancestral karyotype of the Drosophila genus, made up by five pairs of acrocentric chromosomes and one small chromosome pair referred to as “the dot” (Powell 1997, Schaeffer et al. 2008). Although the karyotype has evolved between Drosophila species, polymorphisms in karyotype within a species have not been discovered. In our 2020 work (Flynn et al. 2020) describing inter and intra species satellite DNA variation, we found that one strain in particular, vir00 (15010-1051.00) contained 15% higher satellite DNA abundance than other strains. The elevated satellite DNA abundance in vir00 prompted us to study it further, which we will present here.

Evolution of the sex chromosome arrangement has received abundant attention in both empirical and theoretical studies. Sex chromosome evolutionary studies have mainly made use of sex chromosomes that arose in different time periods (Charlesworth and Charlesworth 2000). In Drosophila, when an autosome fuses to either an X or Y chromosome, a so-called neo-Y chromosome is formed. Because either the fused or unfused version of the chromosome will only be present in males and male Drosophila do not undergo recombination, mutations immediately begin to accumulate through Hill-Robertson interference and other linked-selection processes (Charlesworth and Charlesworth 2000). In the genus Drosophila, autosomes have fused to sex chromosomes multiple independent times (Nozawa *et al.* 2021): *D. pseudoobscura* (10 million years), *D. miranda* (1 million years; Bachtrog 2013), *D. albomicans* (0.24 million years; Wei and Bachtrog 2019), and *D. americana* (29 thousand years; Vieira *et al.* 2006). Neo-sex chromosomes formed by *de novo* sex chromosome fusions are rare and actually under-represented compared to autosomal fusions in *Drosophila* (Anderson *et al.* 2020), and have never been discovered at their infancy before detectable divergence has occurred. How new sex chromosome fusions become stable, and how they compete with the ancestral karyotype within a species is unknown.

Here, we report the discovery of three independent and extremely recent sex-chromosome to autosome fusions in one *D. virilis* strain, vir00. Two fusions involved the X chromosome and one involved the Y chromosome. We isolated four different versions of the same strain that differ in their karyotype (and satellite DNA abundance). We hypothesize that this strain has been more prone to demonstrating chromosome fusions, because of an increased risk of any or multiple of the steps that must occur to detect a fusion. Our findings suggest satellite DNA abundance is a factor that could affect the occurrence of DNA breakage events under stress, the first step in chromosome fusions.

## RESULTS

### Three novel sex chromosome Robertsonian translocations in *D. virilis* strain vir00

In summer 2019, we performed DNA fluorescence in-situ hybridization (FISH) on larval neuroblast nuclei of the vir00 strain that we had obtained in Fall 2018 from the National Drosophila species Stock Center. We discovered that it contained a Y-autosome fusion (Figure 1A). The Y chromosome is recognizable in *D. virilis* because it has a distinct DAPI staining intensity pattern, and contains a distinct arrangement of satellite DNA (Flynn *et al.* 2020). However, all other chromosomes are difficult to distinguish in metaphase spreads, so we could not immediately determine the fusion partner. We observed that the fused chromosome contained the same centromere-proximal satellite as the Y chromosome, AAACTAT (staining faintly with DAPI). To determine if this chromosome fusion was present in other strains from similar geographical locations, we imaged larvae from strains vir08, vir86, and vir48, which were collected from different localities in California and Mexico (vir00 was collected from California). All larvae screened from these other strains contained the same karyotype as the *D. virilis* genome strain (Flynn et al. 2020). We designate the vir00 substrain with the Y fusion as vir00-Yfus.

**Figure 1.**
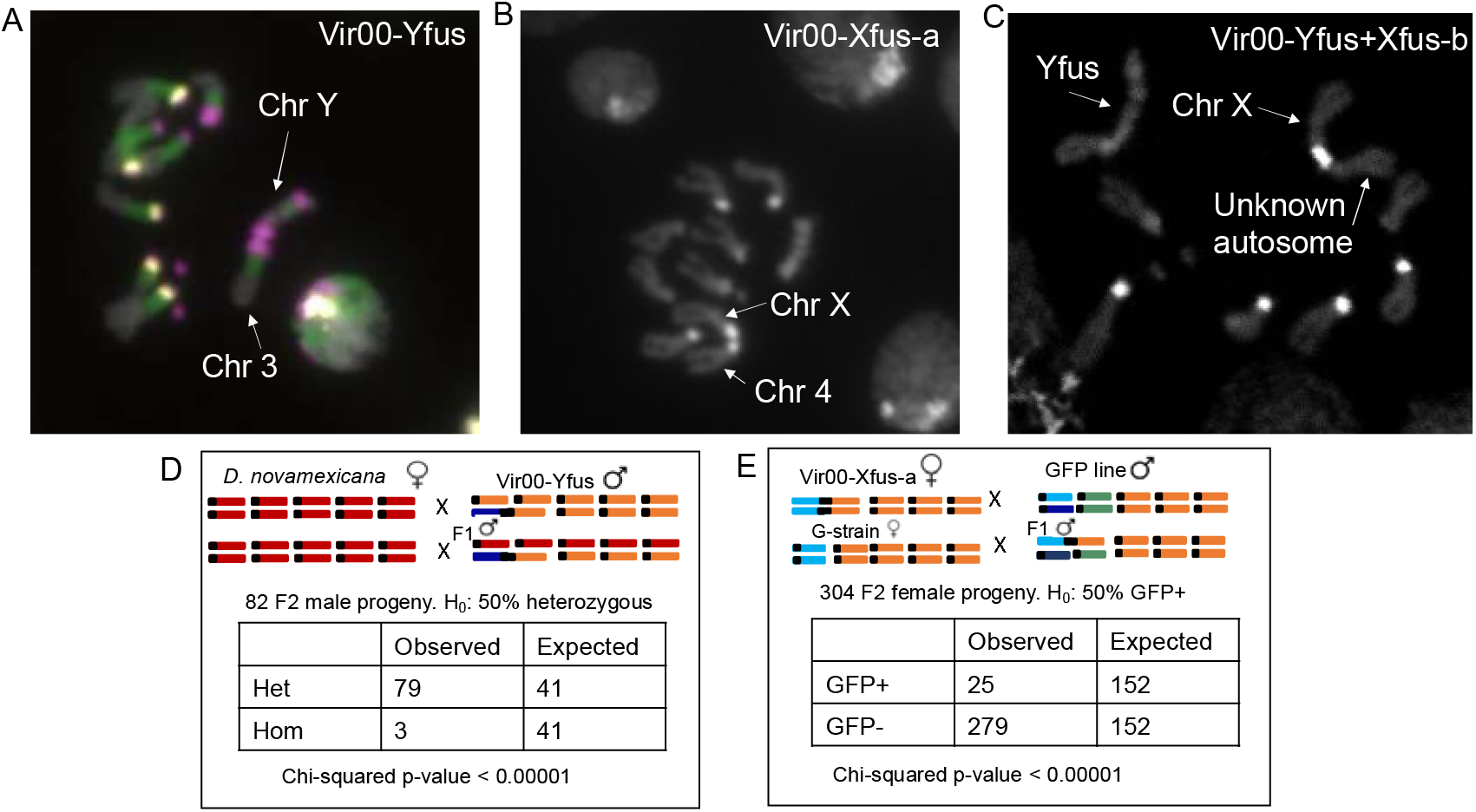
Discovery of three independent fusion events in vir00, and genetic validation of two of them. All metaphase images are of male larval neuropblasts. A) DNA-FISH image demonstrating the Y fusion in vir00-Yfus. b) DAPI staining image demonstrating a X fusion in vir00-Xfus-a. C) DAPI staining image demonstrating a different X fusion in vir00-Yfus+Xfus-b. D) Two-generation crossing experiment to validate of the Y-3 fusion in vir00-Yfus and E) the X-4 fusion in vir00-Xfus-a. Red chromosomes: *D. novamexicana*; orange: wildtype *D. virilis* autosomes; green: autosome containing a GFP marker; light blue: X chromosome; dark blue: Y chromosome.

In Fall 2019, we obtained a second copy of vir00 from the National Drosophila Species Stock Center. We set up single pair crosses and performed larval neuroblast squashes to karyotype multiple male progeny of each cross. Surprisingly, we found that the Y-autosome fusion was not present in any of the larvae karyotyped. However, 3/10 crosses karyotyped contained a different fusion (Figure 1B). This new fusion contained the bright-dapi-staining AAATTAC satellite of both ancestral centromeres, indicating that it is completely distinct from the Y fusion and involved different chromosomes. We hypothesized that this new fusion involved the X chromosome for several reasons; 1) based on centromere satellite identity, it had a 67% probability (Flynn *et al.* 2020); 2) it was found only in single copy in male larvae, but sometimes two copies in female larvae; 3) we observed karyotypes arising from X-Y nondisjunction events in most crosses from this line, such as an XXYY female (Figure S1). We performed single-pair crosses and screened the resulting progeny until we isolated a substrain fixed for the X fusion, and called this substrain vir00-Xfus-a. We retained the stock that contained vir00-Xfus-a segregating with a no-fusion karyotype (vir00-Nofus/vir00-Xfus-a). Additionally, we later isolated a substrain fixed for the no fusion karyotype, vir00-Nofus, by screening single-pair crosses as described above.

In Fall 2021, we obtained a third copy of the vir00 stock from a colleague who had obtained it from the stock center circa 2016 (Yasir Ahmed-Braimah, personal communication). The original purpose of checking this stock copy was to aid in inferring the timeline of the chromosome fusion events, specifically whether the Y fusion was ancestral in the vir00 lineage. We confirmed that this stock contained the above-described Y fusion, but it also had an additional and different fusion segregating, present in about 50% of the larvae imaged (Figure 1C). The new fusion contained the bright-staining AAATTAC satellite on only one of the two ancestral Robertsonian centromeres, in contrast to the other two fusions we previously identified. We performed single-pair crosses and screened the resulting progeny until we isolated a substrain fixed for this new fusion. We also inferred that this fusion involved the X chromosome as well, since once fixed, there were always two copies of the fusion in females (Figure S2), and one copy in males (Figure 1C). Therefore, we designate this substrain as vir00-Yfus+Xfus-b. We inferred that all three fusions we discovered likely represent canonical Robertsonian translocations, in which two acrocentric chromosomes that underwent DNA breakage were fused together at the centromere during repair. We did further experiments on multiple versions of vir00, summarized in Table S1.

### Genetic validation revealed Y-3 and X-4 fusions

We designed two separate two-generation crossing experiments to validate the initially-discovered fusions (vir00-Yfus and vir00-Xfus-a) and identify the autosome each sex chromosome is fused to. Both experiments exploited autosomal markers markers which we could determine if they were segregating non-independently of sex. The first experiment to validate the Y-autosome fusion used crosses between vir00-Yfus and *D. novamexicana* and scoring of polymorphic microsatellite loci on each candidate autosome. We scored 82 F2 male progeny for the Chr3 marker, and 79/82 contained both alleles, whereas the null hypothesis was 50% should contain both alleles (Figure 1D, Figure S3). We determined that the three progeny that did not contain both alleles (Figure 1D Observed-Homozygous) did not contain a Y chromosome because of a nondisjunction event (Figure S4). We concluded that the Y chromosome is fused to Chr3. We also inferred that the Y-3 is fixed in this substrain since no Y chromosomes in the 82 male progeny we assayed segregated independent of Chr3. The other markers on Chr2, Chr4, and Chr5 segregated independently of sex and acted as negative controls (Table S2). We also did a negative control with the same crossing scheme except with vir08 instead of vir00 (Chr3 chi-square p = 0.39, N=22, Table S2).

The second experiment validated the X-autosome fusion in vir00-Xfus-a by crossing the substrain to *D. virilis* transgenic lines containing eye-expressing GFP markers on one of the candidate autosomes. For the crosses to vir95 (which contained a GFP marker inserted on Chr4), we phenotype 304 F2 female progeny, and found that progeny containing a GFP signal were significantly depleted compared to the Mendelian expectation of 50% (Figure 1E). This indicated the X chromosome is fused to Chr4. The same crossing scheme against two other strains containing the GFP marker on Chr2 and Chr5 did not show association of GFP signal with sex (Chi-square p-value > 0.1, Table S3). We also performed a negative control with the *D. virilis* genome strain instead of vir00-Xfus-a crossed to the vir95 line and found the GFP signal was independent of sex (Chi-square p-value = 0.92, Table S3).

We validated that the three fusions are the same genetic line and not the result of contamination from other lines either in our lab or the stock center. We made use of medium-coverage whole genome sequencing from Flynn et al. (2020) to design primers to amplify singleton insertion/deletion variants present only in vir00 (the version that was sequenced was, in hindsight, vir00-Yfus) and not in any other wildtype *D. virilis* strain present in the stock center (Table S4). We designed primers to amplify four loci on chromosomes 2, 3, 5, and 6 which contain a homozygous 12-13 bp deletion in vir00 compared to the reference and the other strains (Table S5). We found that vir00-Yfus, vir00-Xfus-a, vir00-Nofus, and vir00-Yfus+Xfus-b all contained the deletion at each of these loci, supporting that these substrains indeed originated from the same strain vir00 (Figure S5). Although we performed the above indel experiment first, the whole-genome sequencing SNP analysis described in a later section was also concordant with these substrains being the same genetic line.

### Nondisjunction between the X and Y chromosomes is 21% in vir00-Xfus-a

Nondisjunction occurs when homologous chromosomes fail to separate at meiosis, and results in aneuploidy in the progeny. Autosomal and X chromosome aneuploidy is lethal in flies; but Y chromosome aneuploidy is viable: females that have a Y chromosome (XXY) are fertile, males with no Y chromosome (XO) are sterile, and males with two Y chromosomes (XYY) are fertile. Elevated rates of X-Y nondisjunction represent a fitness cost because zygotes with infertile or lethal karyotypes will form at increased frequency. We found some evidence of nondisjunction in the vir00-Yfus genetic validation (3/82 males, Figure 1D, Figure S4), and also common Y chromosome aneuploidy in the stock of vir00-Xfus-a (Figure S1). Since the fusions involve the sex chromosomes, we tested the rate of primary X-Y nondisjunction in males in vir00-Yfus, vir00-Xfus-a, and vir00-Nofus, along with the *D. virilis* genome strain as a control.

We crossed individual males of each strain we tested to genome strain females and genotyped for the presence or absence of the Y chromosome in progeny. Female progeny containing a Y chromosome indicate XY sperm from the father, and male progeny lacking a Y chromosome indicate nullisomic sperm from the father. Since nondisjunction was extremely high in vir00-Xfus-a, 2/7 fathers tested were of XYY karyotype, which we could infer if more than half of his female progeny contained a Y chromosome (Maggert 2014). We eliminated these fathers’ progeny from the primary nondisjunction rate calculation. No fathers tested from other sublines were determined to be XYY. Males without a Y chromosome would not produce progeny. We found that vir00-Yfus had a slightly elevated nondisjunction rate of 4.5%, compared to the genome strain control of 1.2%, but it was not statistically significant with the sample sizes we used (Table 1, p=0.123). vir00-Xfus-a had an extremely high primary nondisjunction rate of 21% (e.g. Figure S6), which was significantly higher than that of all other substrains (p < 0.004, pairwise proportion test, Table 1). Surprisingly, vir00-Nofus had a significantly elevated nondisjunction rate compared to the genome strain control, at 5.7% (p=0.004). This was not statistically different from vir00-Yfus (p=0.231).

**Table 1:** Nondisjunction of the X and Y chromosomes in males is elevated in chromosome fusion lines.

| Line | Nondisjunction rate | Fathers with nondisjunction/total fathers | Aneuploid progeny/ total progeny | Significance group ($p<0.004$) |
| --- | --- | --- | --- | --- |
| GDvir | 1.2% | 1 / 10 | 3 / 248 | a |
| vir00-Yfus | 4.5% | 4 / 7 | 9 / 198 | ab |
| vir00-Nofus | 5.7% | 4 / 6 | 11 / 192 | b |
| vir00-Xfus* | 21% | 5 / 5 | 27 / 127 | c |

\*only including primary nondisjunction

### Satellite DNA abundance has evolved between vir00 substrains

We used Illumina to resequence three vir00 substrains: vir00-Yfus, vir00-Xfus-a, and the mixed stock of vir00-Nofus/Xfus-a. We then used k-Seek to quantify the satellite abundance in each substrain (Wei *et al.* 2014). Satellite abundance was highest in vir00-Yfus compared to vir00-Xfus-a and vir00-Nofus/Xfus-a (Table S6). vir00-Xfus-a had 8% less AAATTAC, which occupies the centromere region of ChrX and Chr4. Overall, vir00-Nofus/Xfus-a contained 12% less pericentromeric satellite DNA compared to vir00-Yfus, with 10% less in pericentromeric AAACTAC, 8% less in AAATTAC, and 13% less in AAACTAT. Given the difference between vir00-Nofus/Xfus-a and the pure vir00-Xfus-a stock, we infer satellite DNA would be even less abundant in vir00-Nofus but unfortunately the X fusion was still segregating with the unfused chromosomes at the time of sequencing.

### Estimated timeline of chromosome fusion events

Since we discovered three different chromosome fusion events in the vir00 stock at different times, we propose here the most likely and parsimonious timeline of events (Figure 2). We used the data we had available, including karyotype, nondisjunction rate, and SNP data from resequencing analysis. Since the Y-3 fusion was fixed in both the pre-2016 (vir00-Yfus+Xfus-b) and 2018 (vir00-Yfus) substrains, we inferred this fusion likely occurred ancestrally. We used resequencing data to estimate how old this fusion may be. The Y-fused version of Chr3 is expected to accumulate mutations independently of the autosomal version of Chr3 over time because of the halt in recombination in male flies. This would be represented by elevated heterozygosity on Chr3 as assayed by aligning short read sequencing data (since reads from both the Y-fused and autosomal copy will map to the same copy of Chr3 in the reference genome). Specifically, if the mutations arose after the fusion of Y-3, they would not be present in any other strains of virilis (assuming no recurrent mutation). Thus, we used our sequencing data for SNP genotyping and found heterozygous singletons unique to vir00-Yfus compared to other non-vir00 strains on each autosome. We found that the number of heterozygous singletons was modestly but significantly enriched on Chr3 in vir00-Yfus: 2.45 SNPs/Mb more than other autosomes (permutation test, p < 0.001). We suggest this elevated density of singleton heterozygous sites may be due to the fusion with the Y chromosome and lack of recombination over several generations. Assuming enrichment was caused by only the Y chromosome and that the neo-Y (Chr3) evolved clonally in a single lineage with single nucleotide mutation rate of 2 x 10^-9^ per nucleotide per generation, we estimate with simulations that the Y-3 fusion occurred 1000-2000 generations ago.

**Figure 2.**
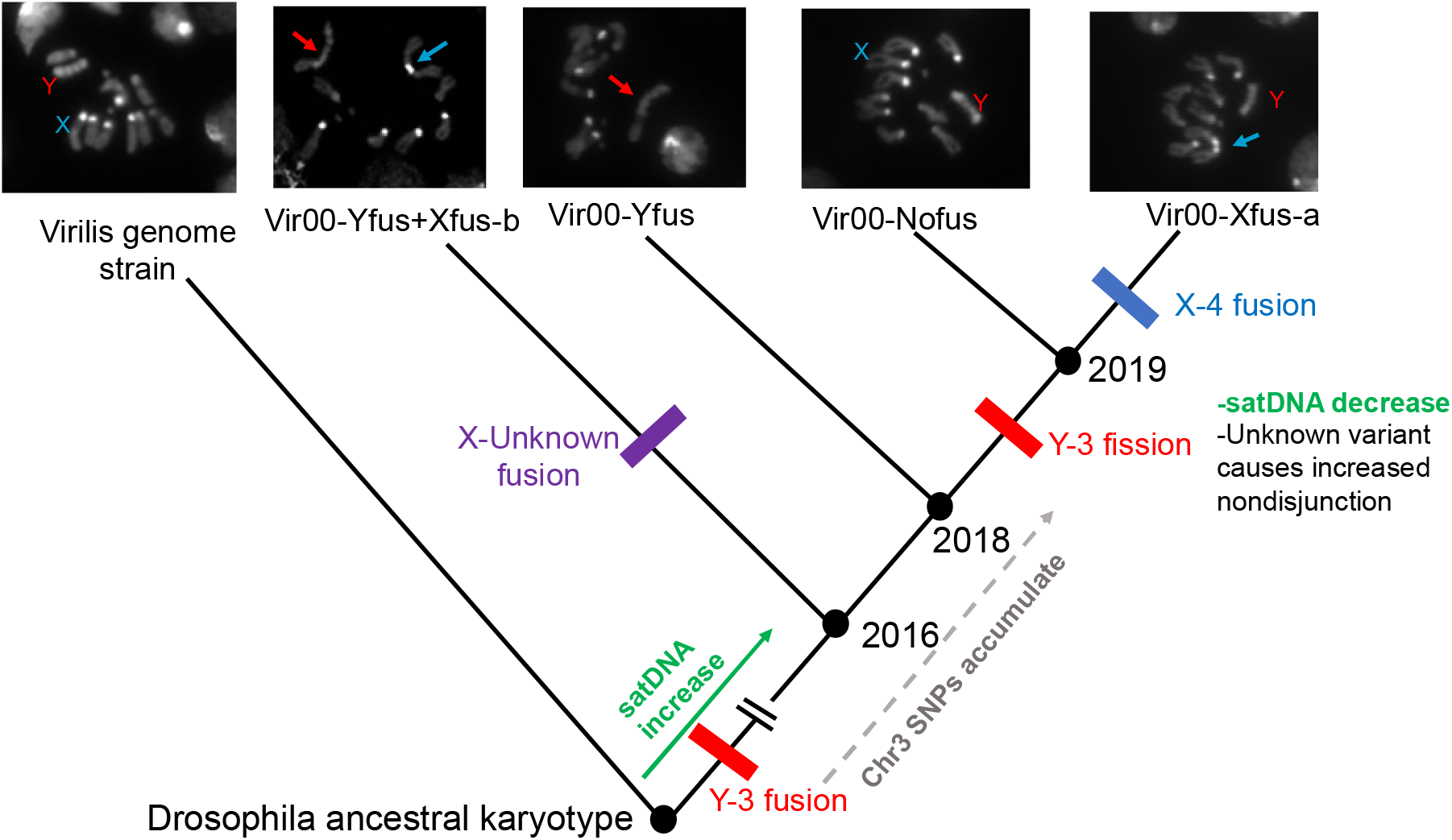
Proposed timeline of chromosome fusion events illustrated on a cladogram (not to scale). The Y-3 fusion, shown by red bars and arrows, appears to be ancestral to the strain and likely occurred 1000-2000 generations ago. The X-Unknown fusion is shown by purple bars and arrows, and the X-4 fusion is shown by blue bars and arrows. Green text describes satellite abundance changes.

Because vir00-Xfus-a was not fixed in the substrain we discovered it in (it was segregating with vir00-Nofus), recombination in heterozygotes would prevent degeneration of the unfused (neo-Y) version of Chr4. Furthermore, we believe the X-4 (Xfus-a) fusion occurred between 2018-2019 since it was not present in the stock we obtained in 2018 or from the pre-2016 stock. We did not observe an enrichment of heterozygous singletons on Chr4 in vir00-Xfus-a. However, there was still a slight enrichment on Chr3 (1.45 SNPs/Mb, permutation test p=0.031), supporting our assumption that the X-4 fusion occurred after the Y-3 fusion broke apart in the same lineage (Figure 2). Furthermore, vir00-Nofus contained a significantly elevated nondisjunction rate of 5.7%, which was surprising since we expected this result only for substrains with chromosome fusions. We think the high nondisjunction rate could be caused by an additional aberration that occurred on the same branch that the Y-3 broke apart. Further studies will be required to test this and determine the nature of the aberration. Although we did not measure the nondisjunction rate in vir00-Yfus+Xfus-b (Table S1), it did not appear as high as vir00-Xfus-a since only one of five crosses we checked had any Y chromosome aneuploidy, supporting its more basal position on the cladogram. An alternative possibility, a tweak to our proposed timeline, is that vir00-Nofus was segregating in the stock undetected since the beginning, and that the Y-3 fusion was lost by drift rather than fission in 2018-2019. Indeed, centromeric fissions are thought to be rare (PláLEK *et al.* 2005). However, we think the long-term segregation of vir00-Yfus and vir00-Nofus is unlikely because the maintenance of fly stocks involves small population sizes which makes it more probable that alleles will be fixed or lost. Our data suggests the Y-3 fusion existed for the longest time, with significant SNPs accumulated on Chr3, making it unlikely that an unfused allele would persist segregating for 1000-2000 generations. Additionally, the Y fusion was completely fixed in both the stock copies we received containing it, whereas both the X fusions were segregating.

### DNA damage levels in response to stress is higher in vir00-Yfus than derived substrains

The unusual commonness of chromosome fusions in vir00 could be caused by an increase in the probability of any (or multiple) of the four steps required to detect a Robertsonian translocation outlined in the introduction. We decided to explore the idea that genetic variation in satellite abundance could contribute generally to the rate of DNA breakage after stress, the fundamental first step that could lead to a Robertsonian translocation or other large-scale rearrangement. We therefore measured DNA damage levels in 7 different *D. virilis* strains, including vir00-Yfus and vir00-Nofus/Xfus-a, with varying abundances of satellite DNA in response to replication stress (induced with the drug gemcitabine) and low-level radiation (10 Gy). For each of seven strains tested, we included a control which was fed with the same liquid food with no gemcitabine and did not receive radiation treatment. We used the comet assay or single-cell gel electrophoresis to measure DNA damage in the male germline in each line, treatment and control (Figure 3). We used the metric “olive moment” to quantify the proportion of damaged DNA per nucleus, which is based on comet shape and DNA staining intensity (Figure S7).

**Figure 3.**
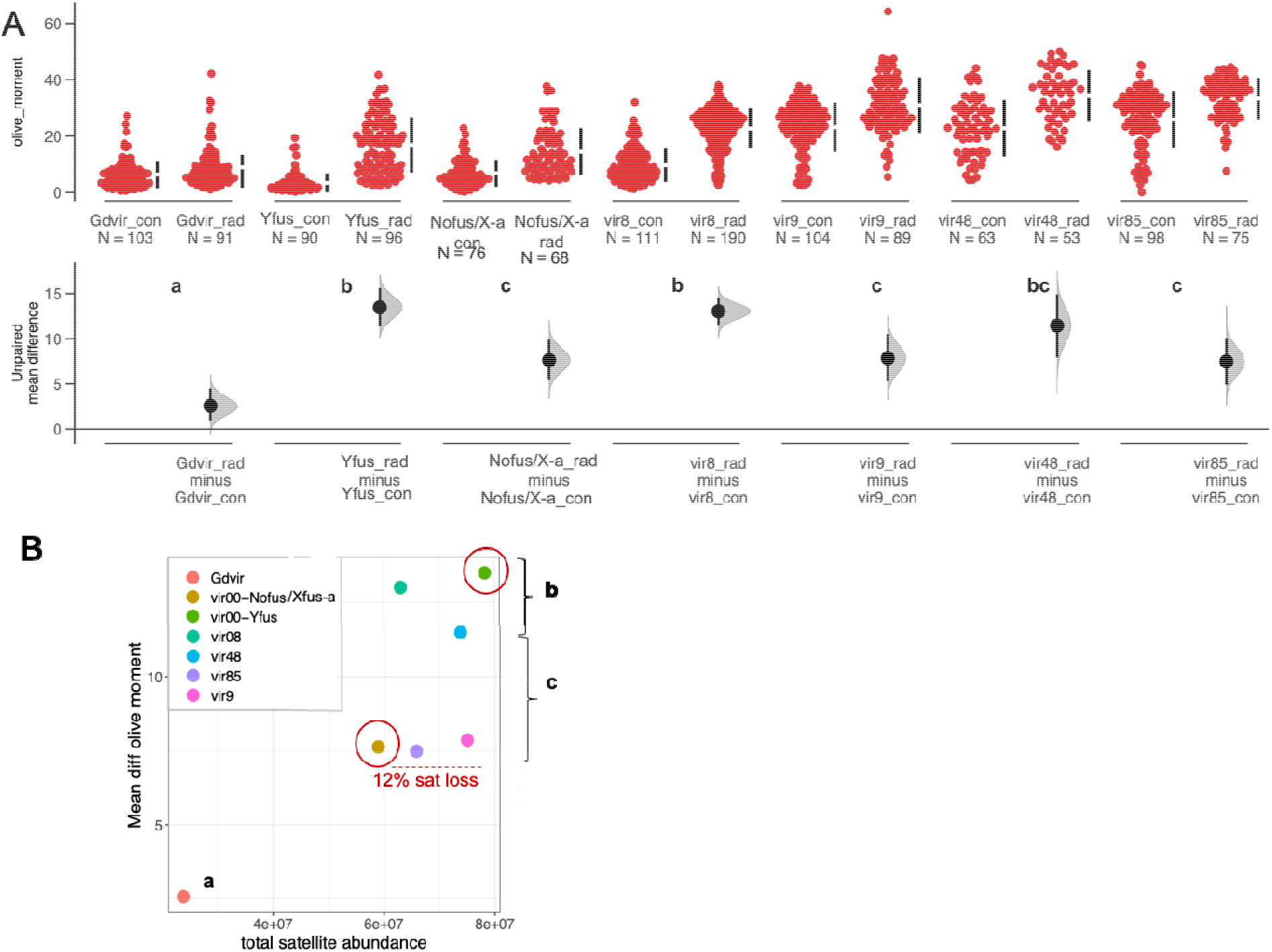
DNA damage levels and satellite abundance in different *D. virilis* strains. A) Olive moment, a statistic of the comet assay, measuring DNA damage for 7 different strains paired with a control and stress treatment. N indicates the number of nuclei analyzed for each treatment. Flies treated with gemcitabine and radiation are suffixed with “rad” and control flies are suffixed with “con”. The lower panel shows the unpaired mean difference between “rad” and “con” for each fly strain (black dot) and 95% confidence intervals (black line) produced from dabestr (5000 bootstrap method). Groups a, b, and c indicate samples with overlapping confidence intervals. B) The satellite DNA abundance of each *D. virilis* strain (x axis) and its unpaired mean difference between treatment and control (y axis). vir00-Nofus/Xfus-a experienced a 12% loss in satellite DNA compared to vir00-Yfus, which is associated with a significantly decreased DNA damage in response to stress.

The strain with the lowest satellite abundance, the genome strain, contained the lowest DNA damage response and this was significantly lower than all other strains tested (Figure 3A). The strain with the highest satellite abundance, vir00-Yfus, had among the highest DNA damage responses, but was not significantly different from two other strains vir8 and vir48 (Figure 3A). Like most other phenotypes, there are likely multiple genetic factors contributing to the variation we found. Ideally, to demonstrate that satellite DNA is a causal factor, we would manipulate satellite DNA abundance and test the DNA damage phenotypes, but multi-megabase long arrays of satellite DNA cannot be manipulated with traditional genome editing. However, vir00-Yfus and vir00-Nofus/Xfus-a are expected to have an almost identical genetic background except for a chromosome fusion difference and a satellite abundance difference. Thus, if there is a difference in DNA damage response between these substrains, it may be caused by differing satellite abundance. vir00-Nofus/Xfus-a, which contained 12% less satellite DNA than vir00-Yfus, had a significantly reduced DNA damage response, concordant with our expectations that satellite DNA plays a role (Figure 3B), likely in addition to other factors.

## DISCUSSION

Here, we report the discovery of a *D. virilis* strain that has undergone an unusual number of chromosome fusion (and likely fission) events very recently. Chromosome rearrangements such as Robertsonian translocations are rare occurrences, but are often the source of karyotype evolution between species (Mayrose and Lysak 2021). In humans, rearrangements resulting from breaks near the centromere are associated with miscarriages and in the case of somatic rearrangements, cancer (Barra and Fachinetti 2018). Previous studies have found repetitive element involvement in chromosome rearrangements in multiple species (Paço *et al.* 2015; Reis *et al.* 2018). Here, we suggest satellite abundance may influence the risk of DNA damage and chromosome rearrangements. There are several possible mechanisms that might cause excess satellite DNA to increase the risk of DNA breakage. Satellite DNA may form complex structures, loops, or non-B DNA, and increasing the length of the arrays may make these regions more unstable in cis (Barra and Fachinetti 2018). It may be challenging for polymerases to replicate many megabases of tandem satellite sequence, and longer arrays may have a higher risk of stalling polymerases and thus DNA breaks in cis (Barra and Fachinetti 2018). Finally, increased satellite DNA may titrate away binding proteins that maintain genome stability, in trans (Francisco and Lemos 2014; Brown *et al.* 2020a; Giunta *et al.* 2021).

Our study has several limitations. Firstly, the timeline of unexpectedly discovering different karyotypes in vir00 at different times resulted in some substrains not having data collected for all experiments (Table S1). For example, vir00-Nofus/Xfus-a was segregating in our sequencing and comet assay experiment, but vir00-Nofus was isolated for the nondisjunction assay. vir00-Yfus+Xfus-b was discovered very recently and only strain validation was able to be completed at this time. Furthermore, our experiment demonstrating a higher damage level between the control and stressed flies in vir00-Yfus may not be directly applicable to the risk of DNA breaks and genome instability in natural conditions. Vir48 and vir08 had a similar olive moment difference to vir00-Yfus, but did not exhibit chromosome fusions, suggesting other factors not studied here playing an important role, possibly at other steps. Although we find a difference in the DNA damage in response to stress between vir00 substrains with different abundances of satellite DNA, their fusion status is also different. We cannot eliminate the possibility that the presence of the Y fusion itself increased the rate of DNA damage instead of the abundance of satellite DNA.

We identified elevated nondisjunction in some vir00 substrains. Surprisingly, the rate of nondisjunction was 5-fold higher in vir00-Xfus compared to vir00-Yfus. With a nondisjunction rate of 21%, a high proportion of abnormal karyotypes are produced, such as XO (sterile), XYY (viable and fertile), and XXY or XXYY (viable and fertile), all of which we found in cytological samples. Further mating between these abnormal karyotypes will produce significant proportions of sterile or inviable karyotypes like XYYY or XXX, which will further decrease the fitness of this line. Furthermore, karyotypes with extra Y chromosomes such as XYY and XXY have been found to have decreased lifespan (Brown *et al.* 2020b). The extreme nondisjunction rate vir00-Xfus-a but not in vir00-Yfus, indicates there may be additional rearrangements on the X and/or Y that occurred during fission of the Y-3 and disrupted proper pairing and segregation of the sex chromosomes. Furthermore, the nondisjunction rate in vir00-Nofus is still significantly higher than the genome strain, despite the same karyotype. We believe this indicates a remaining rearrangement in vir00-Nofus affecting the pairing and/or disjunction of the X-Y. We only found only a modest difference in estimated rDNA copy number between the substrains (Table S8), which has been found to mediate pairing of the X and Y (McKee and Karpen 1990). Detailed analysis of structural rearrangements in the heterochromatin along with cytological analysis will be required to determine the mechanism of the elevated nondisjunction rates.

We have isolated four different versions of the strain vir00, each with a different karyotype: vir00-Nofus, vir00-Xfus-a, vir00-Yfus, vir00-Yfus+Xfus-b, which will be useful for a variety of future studies. The vir00 fusion substrains will be useful for studying centromere identity. In both the Y-3 and X-4 fusions, two spherical regions of satellite DNA are present at the centromere of these fusions, representing one from each acrocentric chromosome (Figure 1). We note that the X-4 fusion in vir00 is homologous to an independent X-4 fusion in *D. americana* 29 thousand years ago, a species only 4.5 million years diverged. In the X-4 fusion of *D. americana,* there is only one discrete region of centromeric satellite (Flynn *et al.* 2020), unlike what we found here. In female meiosis where chromosomes can compete to get into the oocyte rather than the polar body, “stronger” centromeres may have an advantage (Malik and Bayes 2006). A “supercentromere” resulting from a centromere-centromere fusion is one possible way to do this, and in *D. americana* the X-4 fusion has biased transmission into the egg (Stewart *et al.* 2019). Furthermore, all the fusions we discovered involve sex chromosomes and create neo-sex chromosomes, to our knowledge the most recently formed neo-sex chromosomes reported. Broadly, a species is generally assumed to have a fixed karyotype, and only one other exceptional system to our knowledge has been described to have polymorphic chromosome fusions within a species (house mouse, PláLEK *et al.* 2005). How rare the strain vir00 is compared to other strains of other species, and the dynamics and maintenance of divergent karyotypes within a species is a relatively unexplored field.

## METHODS

Scripts required to reproduce the computational results are available here: https://github.com/jmf422/D-virilis-fusion-chromosomes

### Neuroblast squashes and satellite DNA FISH

We dissected brains from wandering 3^rd^-instar larvae and performed the fixation steps as in (Larracuente and Ferree 2015). Specifically, we placed brains in sodium citrate solution for 6 minutes before fixation. After fixation and drying of slides, we applied Vectashield dapi mounting medium. We performed DNA-FISH on vir00-Yfus, which allowed us to confidently identify the Y chromosome based on its unique satellite DNA composition. We used the same fixation and staining protocol as (Flynn *et al.* 2020). We imaged metaphase cells using a 100x oil objective on an Olympus fluorescent microscope and Metamorph capture system at the Cornell Imaging Facility.

### vir00-Yfus Y-autosome fusion validation

We designed an experiment that would both validate the Y chromosome fusion and to distinguish which autosome is fused. We first designed primers flanking microsatellite loci on all four autosomes that met the following criteria: 1) had 100% conserved non-repetitive and unique priming sites between *D. virilis* and *D. novamexicana*; 2) amplicon length differed between the species by at least 15 bp as to be distinguished on an agarose gel; 3) locus contained a mono or tri nucleotide repeat; 4) locus length ~200 bp. We next set up a two-generation crossing scheme (Figure 1A). We crossed *D. novamexicana* virgin females with vir00-Yfus males and selected the male progeny, which we backcrossed to *D. novamexicana* virgin females. We then genotyped the male F2 progeny from this cross at the 4 sets of primers corresponding to the four non-dot autosomes (Chr2, 3, 4, 5) (Table S2). We performed single-fly DNA extraction in strip tubes with Tris-EDTA buffer and 0.2 mg/mL proteinase K. We did 12 uL standard PCR reactions (3 min at 95, 30 cycles of 30 sec 95, 30 sec 55, 50 sec 72, final extension 5 min). Each primer on each PCR plate had a homozygous (*D. novamexicana*) and heterozygous (D. *novamexicana-D.virilis* F1 hydrid) control. We then ran the PCR product on 2.5% agarose gels. If there was indeed an autosome fused to the Y chromosome, we would expect to see 100% of the male progeny being heterozygous for the *virilis* and *novamexicana* alleles (except for rare cases of non-disjunction). For the autosomes that are not fused, we would expect to see 50% of the progeny being homozygous for the *novamexicana* allele, and half heterozygous, due to Mendel’s law of random segregation. We successfully validated the existence of the Y fusion, and found that it is fused to chromosome 3 (Muller D) (Table S2, Figure 1). There were 3 male progeny that were homozygous for the Chr3 *novamexicana* allele. We verified that these were cases of nondisjunction (opposed to the Y fusion not being fixed in this subline) by finding that the Y chromosome was absent in controlled Y chromosome PCR assays (Figure S4).

### Isolation of vir00-Xfus-a and vir00-Nofus

The first X fusion was found to be segregating with a no fusion substrain in the 2019 stock of vir00. We wanted to isolate these into two separate substrains where the karyotype is fixed. From the progeny of the three original crosses in which we found the X fusion, we made 10 single pair crosses and did neuroblast squashes of 6-8 larval progeny per cross, including both sexes. By chance, we should be able to find a cross in which the mother had two copies of the fusion and the father had a single copy – in which the derived line would be fixed for the fusion. If all progeny imaged contained the fusion (and females contained two copies of the fusion), then it is likely that this was the case. We created this line, and call it vir00-Xfus-a. We maintained a line isolated from the 2019 stock that had low frequency of the X fusion (~25%), and called it vir00-Nofus/Xfus-a. We used this strain for several experiments (Table S1). By the time we completed the nondisjunction assays, we had isolated a pure Nofus substrain (vir00-Nofus, Table S1), so we only performed nondisjunction assays on fixed versions of the vir00 substrains.

### vir00-Xfus-a X-autosome fusion validation

We obtained transgenic strains with GFP (or Blue) insertions which are expressed in the eye and larval brain from the National Drosophila Species Stock Center (vir95, vir121, vir117). Stern *et al.* (2017) found the insertion sites of these lines. We chose lines which contained the GFP marker on candidate autosomes Chr2, Chr4, and Chr5. Chr3 was not a candidate because it is fused to the Y in vir00-Yfus and contains a different centromeric satellite. Before setting up crosses, we screened 10-20 larvae of each line with a fluorescent microscope to ensure the transgene had not drifted to low frequency. Larvae containing the transgene demonstrated the GFP signal in their brain. We chose to phenotype at the larval stage since we would be crossing GFP strains to wildtype red-eyed flies and the visibility of GFP in the adult eye would be low. We then designed a crossing scheme which would allow us to both validate that the X chromosome was fused and distinguish which autosome it was fused to (Figure 1B). We crossed GFP-line males to vir00-Xfus virgin females. We then selected the male F1 progeny and crossed them to virilis genome strain virgin females. We then phenotyped F2 larvae, classifying each as either GFP positive or negative. When the phenotyped flies emerged, we sexed and counted them. If the candidate autosome is fused to the X, we would expect sex to segregate with the GFP marker: all female progeny will be GFP negative, and all male progeny will be GFP positive (except for phenotyping errors or rare nondisjunction events). For all other lines, sex should not segregate with GFP status. We performed negative control crosses in which the parental cross was replaced by genome strain virgin females, to ensure the crossing scheme produced the expected results (Table S3). We found that the X chromosome is fused to Chr4 (Muller B).

### Validation of the three versions of vir00 with private fixed indels

We first re-sequenced the same sequencing libraries produced from Flynn et al. (2020) to obtain high enough coverage for genotyping. Raw data, including the original sequencing and the resequencing are available at NCBI SRA PRJNA548201. We then used GATK recommended practices to do genotyping of our low-coverage whole genome sequencing data. We used vcftools to subset singletons present only in vir00, which was, in hindsight, vir00-Yfus. We then used GATK’s SelectVariants to select only non-reference homozygous indels 12 bp or more with a depth of at least 10 in vir00 and at least 2 in the other strains. We then manually inspected each potential candidate in IGV to ensure: no reads in other strains supported the indel, all reads in vir00 supported the indel, and there were no nearby indels in other strains. We then designed primers for the four loci (on Chr 2, 3, 5, 6) that met these criteria and also had enough SNP-free sequence flanking the indel in order to design primers that would amplify a locus 100-200 bp equally in all strains (Table S5). We performed PCR and gel electrophoresis (2.5% gel, 98 V, 90 min).

### DNA damage assays

We chose two stressors that would moderately increase the rate of DNA breaks and allow us to potentially detect differences between strains. Gemcitabine is a nucleoside analog that induces replication stress by stalling polymerases, and also sensitizes cells to radiation via the RAD51 pathway (Kobashigawa *et al.* 2015). We selected dosage and a fly-feeding regime based on (Kislukhin *et al.* 2012). Ionizing radiation has long been used to increase the rates of DNA breaks in flies for mutagenesis. We chose a dose ¼ - ½ of what has been typically used in mutagenesis(Carlson and Southin 1962).

We collected male flies 0-1 days old and fed them gemcitabine (0.718 mM) mixed with liquid food in vials with 8-12 adult flies as in (Kislukhin *et al.* 2012). Liquid food consisted of 12.5g sucrose, 17.5 g dry yeast, 5 mL corn syrup, and 95 mL PBS (autoclaved for 30 min immediately after adding the yeast). Flies were fed the drug for 7-9 days before radiation. Controls were fed with the same liquid food for 7-9 days, except no gemcitabine was added. Flies were moved to fresh vials every 3-4 days. For radiation treatment, we transferred flies into 50 mL conical tubes with 5 mL agar because these tubes were compatible with the radiation source. Control flies were also transferred to new tubes. We used a J.L. Shepherd & Associates Mark I Irradiator with 1,100 Ci of Cs-137, and flies were irradiated at approximately 400 rad/min for a total of 10 Gy. In one case, for the GDvir stress treatment, the radiation was not stopped on time so 4 extra Gy were applied. We believe this did not affect our results, especially because the GDvir strain had the lowest DNA damage increase with gemcitabine and radiation stress. We did comet assays to measure DNA damage (Angelis *et al.* 1999) over three different dates (Table S7), but ensured experimental conditions were practically identical each time. For some samples we had to combine results from two different dates to have enough nuclei for statistical analysis (Table S7). We dissected testes from approximately 8 flies from each treatment within one hour of radiation treatment to minimize the opportunity for breakage repair (Shetty *et al.* 2017). We then homogenized the testes tissue using a dounce, filtered the homogenate through a 40 micron filter to remove debris, and centrifuged and resuspended the cells to approximately 10^5^ cells/mL.

We next performed the alkaline comet assay as directed by the Enzo comet kit (ADI-900-166), which provides higher sensitivity than the neutral comet assay (Angelis *et al.* 1999). We imaged slides on a metamorph imaging system at 10x magnification using a fluorescent green filter to detect the CyGreen dye included in the comet kit. We quantified damage levels using the software OpenComet as a plugin in ImageJ (Gyori *et al.* 2014). We filtered called nuclei that were not comet shapes or contained background interference. We used the measure of “olive moment,” which is the product of the percent of DNA in the tail and distance between intensity-weighted centroids of head and tail (Gyori *et al.* 2014) as the statistic to compare between strains and treatments.

### Resequencing sublines to determine differences in their satellite abundance

Pools of 6 male flies were DNA extracted with Qiagen DNeasy blood and tissue kit. PCR-free libraries were then prepared with Illumina TruSeq PCR-free library prep. Libraries were sequenced on a NextSeq 500 single end 150 bp. We removed adapters and poly-G signal with fastp and then ran k-Seek to count satellite abundances (Wei *et al.* 2014). We used average read depth to normalize the kmer counts. We also mapped the reads to the *D. virilis* rDNA consensus sequence (http://blogs.rochester.edu/EickbushLab/?page_id=602) to estimate the rDNA copy number in the three vir00 substrains as well as a vir08 as a control (Table S4, S8).

### Using sequencing data to estimate the age of the Y-3 fusion

Scripts for this section are available here: https://github.com/jmf422/D-virilis-fusion-chromosomes/tree/main/simulate_degradation. We used the sequencing data from Flynn et al. (2020) and the resequencing of the same strains, in addition to data produced here for vir00-Yfus, vir00-Xfus-a, and vir00-Nofus/Xfus-a and 10 other *D. virilis* strains. All raw data is available at NCBI SRA PRJNA548201. We mapped the data to the RS2 genome assembly using bowtie2. We then genotyped with GATK following standard procedures (McKenna *et al.* 2010). We extracted heterozygous singleton sites for vir00, and counted how many occurred on each autosome. We calculated the enrichment on Chr3 in vir00-Yfus based on the difference from the average SNP density on the other autosomes (excluding the dot chromosome Chr6). To determine whether this enrichment of SNPs was significant based on the size of the chromosome and the number of mutations, we randomly permuted the total number of heterozygous singleton SNPs on all autosomes and calculated the proportion falling on Chr3, and repeated this 1000 times.

We then performed simple simulations to determine approximately how many generations of mutation accumulation without recombination or selection would result in the enrichment we observed. Since heterozygous singletons are challenging for the genotyper to call with moderate coverage sequencing data, we incorporated this into our simulation. First, we made the genome assembly diploid then used mutation-simulator (Kühl *et al.* 2020) to simulate random mutations (transition/transversion ratio 2.0) at a rate of 2 x 10^-9^ per bp per generation for 500, 1000, 2000, and 5000 generations on one copy of Chr3 only. We then simulated Illumina reads with ART (Huang *et al.* 2012) at the same depth as we have for vir00-Yfus in our real data (23 x haploid or 11.5 x diploid). We next used standard GATK genotyping and selected out heterozygous singletons on Chr3. We repeated the simulation 10 times for each number of generations to get a range of values. The empirical enrichment fell in between what we found in the simulations for 1000 and 2000 generations.

### Nondisjunction assays

We crossed a single male from vir00-Yfus, vir00-Xfus-a, vir00-Nofus (fixed), and GDvir (control) to one or two GDvir (genome strain vir87) females. We collected the virgin progeny from each cross, extracted DNA with a squish-proteinase K prep, and genotyped with PCR and gel electrophoresis for the presence or absence of the Y chromosome in up to 16 female and 16 male progeny. We amplified a locus unique to the Y chromosome (primers designed by Yasir Ahmed-Braimah for a different project, Table S5). For a subset of individuals, we also performed multiplex controls with an autosomal locus. Otherwise, we performed DNA extractions in large batches with the same proteinase K mixture to minimize the chance of DNA extraction failure. A very small quantity of DNA is required for a standard PCR with robust primers. To control for the completeness of the PCR mastermix, we included male samples in the same batch as female samples. If a male lacked a Y chromosome, we inferred the father’s sperm was missing the Y chromosome (nullisomic), and if a female contained a Y chromosome, we inferred the father’s sperm contained both X and Y. We used R prop.test to evaluate whether there were any differences between nondisjunction proportions for the different strains. After finding this highly significant, we used pairwise.prop.test in R with Holm-Bonferroni multiple test correction to determine which pairs of substrain nondisjunction rates were significantly different from each other.

## Acknowledgements

We thank Yasir Ahmed-Braimah for discussions and use of Y chromosome primers. We greatly appreciate the use of the Cs-137 irradiator from the Robert Weiss lab and to Amanda Loehr for operating the device. Asha Jain prepared sequencing libraries for this project, and Yassi Hafezi provided advice on the nondisjunction assays. Fluorescent images were taken at Cornell Imaging facility. This work was supported by National Institutes of Health Grant number GM119125 to A.G.C. and Daniel Barbash. We also thank members of the Clark lab for discussions and encouragement on this project.

## References

Anderson N. W., C. E. Hjelmen, and H. Blackmon, 2020 The probability of fusions joining sex chromosomes and autosomes. Biol. Lett. 16: 20200648.

Angelis K. J., M. Dusinská, and A. R. Collins, 1999 Single cell gel electrophoresis: detection of DNA damage at different levels of sensitivity. Electrophoresis 20: 2133–2138.

Bachtrog D., 2013 Y-chromosome evolution: emerging insights into processes of Y-chromosome degeneration. Nat. Rev. Genet. 14:113–124.

Balzano E., F. Pelliccia, and S. Giunta, 2020 Genome (in)stability at tandem repeats. Semin. Cell Dev. Biol. https://doi.org/10.1016/j.semcdb.2020.10.003

Barra V., and D. Fachinetti, 2018 The dark side of centromeres: types, causes and consequences of structural abnormalities implicating centromeric DNA. Nat. Commun. 9: 4340.

Bilinski P., P. S. Albert, J. J. Berg, J. A. Birchler, M. N. Grote, et al., 2018 Parallel altitudinal clines reveal trends in adaptive evolution of genome size in Zea mays. PLoS Genet. 14: e1007162.

Black E. M., and S. Giunta, 2018 Repetitive Fragile Sites: Centromere Satellite DNA As a Source of Genome Instability in Human Diseases. Genes 9. https://doi.org/10.3390/genes9120615

Braekeleer M. D., and T.-N. Dao, 1990 Cytogenetic studies in couples experiencing repeated pregnancy losses. Hum. Reprod. 5: 519–528.

Brown E. J., A. H. Nguyen, and D. Bachtrog, 2020a The Drosophila Y Chromosome Affects Heterochromatin Integrity Genome-Wide. Mol. Biol. Evol. 37: 2808–2824.

Brown E. J., A. H. Nguyen, and D. Bachtrog, 2020b The Y chromosome may contribute to sex-specific ageing in Drosophila. Nat Ecol Evol 4: 853–862.

Carlson E. A., and J. L. Southin, 1962 Comparative mutagenesis of the dumpy locus in Drosophila melanogaster. I. X-ray treatment of mature sperm--frequency and distribution. Genetics 47: 321–336.

Cechova M., R. S. Harris, M. Tomaszkiewicz, B. Arbeithuber, F. Chiaromonte, et al., 2019 High satellite repeat turnover in great apes studied with short-and long-read technologies. Mol. Biol. Evol. https://doi.org/10.1093/molbev/msz156

Charlesworth B., C. H. Langley, and W. Stephan, 1986 The evolution of restricted recombination and the accumulation of repeated DNA sequences. Genetics 112: 947–962.

Charlesworth B., and D. Charlesworth, 2000 The degeneration of Y chromosomes. Philosophical Transactions of the Royal Society of London. Series B: Biological Sciences 355:1563–1572.

Featherstone C., and S. P. Jackson, 1999 DNA double-strand break repair. Curr. Biol. 9: R759–R761.

Ferree P. M., and D. A. Barbash, 2009 Species-specific heterochromatin prevents mitotic chromosome segregation to cause hybrid lethality in Drosophila. PLoS Biol. 7: e1000234.

Flynn J. M., M. Long, R. A. Wing, and A. G. Clark, 2020 Evolutionary Dynamics of Abundant 7-bp Satellites in the Genome of Drosophila virilis. Mol. Biol. Evol. 37:1362–1375.

Francisco F. O., and B. Lemos, 2014 How Do Y-Chromosomes Modulate Genome-Wide Epigenetic States: Genome Folding, Chromatin Sinks, and Gene Expression. Journal of Genomics 2: 94–103.

Fry K., and W. Salser, 1977 Nucleotide sequences of HS-alpha satellite DNA from kangaroo rat Dipodomys ordii and characterization of similar sequences in other rodents. Cell 12:1069–1084.

Gall J., E. Cohen, and M. Polan, 1971 Repetitive DNA sequences in Drosophila. Chromosoma 33. https://doi.org/10.1007/bf00284948

Gall J. G., and D. D. Atherton, 1974a Satellite DNA sequences in Drosophila virilis. Journal of Molecular Biology 85: 633–664.

Gall J. G., and D. D. Atherton, 1974b Satellite DNA sequences in Drosophila virilis. J. Mol. Biol. 85: 633–664.

Giunta S., S. Hervé, R. R. White, T. Wilhelm, M. Dumont, et al., 2021 CENP-A chromatin prevents replication stress at centromeres to avoid structural aneuploidy. Proc. Natl. Acad. Sci. U. S. A. 118. https://doi.org/10.1073/pnas.2015634118

Gyori B. M., G. Venkatachalam, P. S. Thiagarajan, D. Hsu, and M.-V. Clement, 2014 OpenComet: an automated tool for comet assay image analysis. Redox Biol 2: 457–465.

Huang W., L. Li, J. R. Myers, and G. T. Marth, 2012 ART: a next-generation sequencing read simulator. Bioinformatics 28: 593–594.

Jagannathan M., and Y. M. Yamashita, 2021 Defective satellite DNA clustering into chromocenters underlies hybrid incompatibility in Drosophila. bioRxiv 2021.04.16.440167.

Kislukhin G., M. L. Murphy, M. Jafari, and A. D. Long, 2012 Chemotherapy-induced toxicity is highly heritable in Drosophila melanogaster. Pharmacogenet. Genomics 22: 285–289.

Kobashigawa S., K. Morikawa, H. Mori, and G. Kashino, 2015 Gemcitabine Induces Radiosensitization Through Inhibition of RAD51-dependent Repair for DNA Double-strand Breaks. Anticancer Res. 35: 2731–2737.

Kühl M. A., B. Stich, and D. C. Ries, 2020 Mutation-Simulator: Fine-grained simulation of random mutations in any genome. Bioinformatics. https://doi.org/10.1093/bioinformatics/btaa716

Larracuente A. M., and P. M. Ferree, 2015 Simple method for fluorescence DNA in situ hybridization to squashed chromosomes. J. Vis. Exp. 52288.

Maggert K. A., 2014 Reduced rDNA Copy Number Does Not Affect “Competitive” Chromosome Pairing in XYY Males of Drosophila melanogaster. G3 Genes | Genomes | Genetics 4: 497–507.

Mayrose I., and M. A. Lysak, 2021 The Evolution of Chromosome Numbers: Mechanistic Models and Experimental Approaches. Genome Biol. Evol. 13. https://doi.org/10.1093/gbe/evaa220

McKee B. D., and G. H. Karpen, 1990 Drosophila ribosomal RNA genes function as an X-Y pairing site during male meiosis. Cell 61: 61–72.

McKenna A., M. Hanna, E. Banks, A. Sivachenko, K. Cibulskis, et al., 2010 The Genome Analysis Toolkit: a MapReduce framework for analyzing next-generation DNA sequencing data. Genome Res. 20:1297–1303.

Miga K. H., Y. Newton, M. Jain, N. Altemose, H. F. Willard, et al., 2014 Centromere reference models for human chromosomes X and Y satellite arrays. Genome Res. 24: 697–707.

Nozawa M., Y. Minakuchi, K. Satomura, S. Kondo, A. Toyoda, et al., 2021 Evolutionary trajectories of three independent neo-sex chromosomes in Drosophila. Cold Spring Harbor Laboratory 2021.03.11.435033.

Paço A., F. Adega, N. Meštrović, M. Plohl, and R. Chaves, 2015 The puzzling character of repetitive DNA in Phodopus genomes (Cricetidae, Rodentia). Chromosome Res. 23: 427–440.

Petitpierre E., C. Juan, J. Pons, M. Plohl, and D. Ugarkovic, 1995 Satellite DNA and constitutive heterochromatin in tenebrionid beetles, pp. 351–362 in Kew Chromosome Conference IV. London: Royal Botanic Gardens,.

PláLEK J., H. C. Hauffe, and J. B. Searle, 2005 Chromosomal variation in the house mouse. Biological Journal of the Linnean Society 84: 535–563.

Reis M., C. P. Vieira, R. Lata, N. Posnien, and J. Vieira, 2018 Origin and Consequences of Chromosomal Inversions in the virilis Group of Drosophila. Genome Biol. Evol. 10: 3152–3166.

Schulz R., L. A. Underkoffler, J. N. Collins, and R. J. Oakey, 2006 Nondisjunction and transmission ratio distortion ofChromosome 2 in a (2.8) Robertsonian translocation mouse strain. Mamm. Genome 17: 239–247.

Shetty V., N. J. Shetty, S. R. Ananthanarayana, S. K. Jha, and R. C. Chaubey, 2017 Evaluation of gamma radiation-induced DNA damage in Aedes aegypti using the comet assay. Toxicol. Ind. Health 33: 930–937.

Stern D. L., J. Crocker, Y. Ding, N. Frankel, G. Kappes, et al., 2017 Genetic and Transgenic Reagents for Drosophila

Stewart N. B., Y. H. Ahmed-Braimah, D. G. Cerne, and B. F. McAllister, 2019 Female meiotic drive preferentially segregates derived metacentric chromosomes in Drosophila. bioRxiv 638684.

Subirana J. A., M. M. Albà, and X. Messeguer, 2015 High evolutionary turnover of satellite families in Caenorhabditis. BMC Evol. Biol. 15: 218.

Thakur J., J. Packiaraj, and S. Henikoff, 2021 Sequence, Chromatin and Evolution of Satellite DNA. Int. J. Mol. Sci. 22. https://doi.org/10.3390/ijms22094309

Vieira C. P., A. Almeida, J. D. Dias, and J. Vieira, 2006 On the location of the gene(s) harbouring the advantageous variant that maintains the X/4 fusion of Drosophila americana. Genet. Res. 87:163–174.

Wei K. H.-C., J. K. Grenier, D. A. Barbash, and A. G. Clark, 2014 Correlated variation and population differentiation in satellite DNA abundance among lines of Drosophila melanogaster. Proc. Natl. Acad. Sci. U. S. A. 111: 18793–18798.

Wei K. H.-C., S. E. Lower, I. V. Caldas, T. J. S. Sless, D. A. Barbash, et al., 2018 Variable Rates of Simple Satellite Gains across the Drosophila Phylogeny. Mol. Biol. Evol. 35: 925–941.

Wei K. H.-C., and D. Bachtrog, 2019 Male recombination produced multiple geographically restricted neo-Y chromosome haplotypes of varying ages that correlate with onset of neo-Y decay in Drosophila albomicans. PLoS Genet. 15:e1008502.

