## Supplemental figures for "Three recent sex chromosome-to-autosome fusions in a *Drosophila virilis* strain with high satellite content"

**Supplementary Figures**

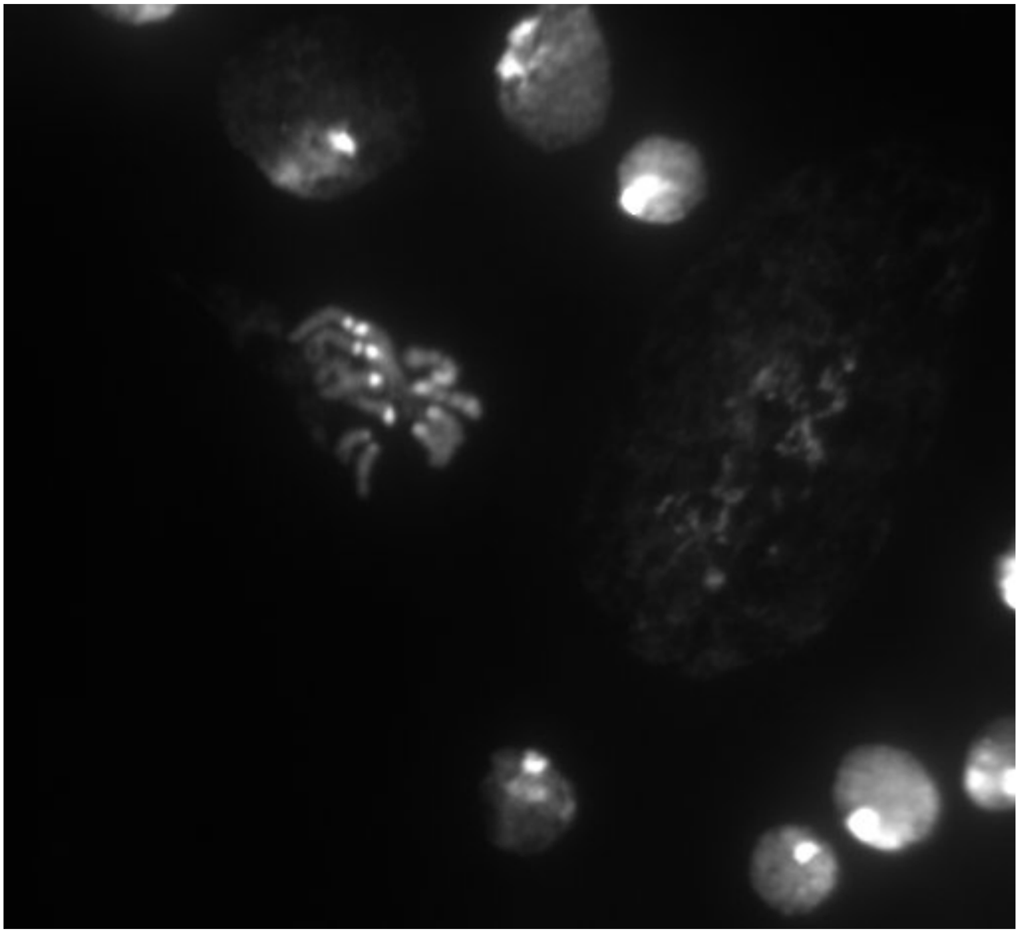

**Figure S1**. XXYY karyotype in vir00-Xfus female larva neuroblast.

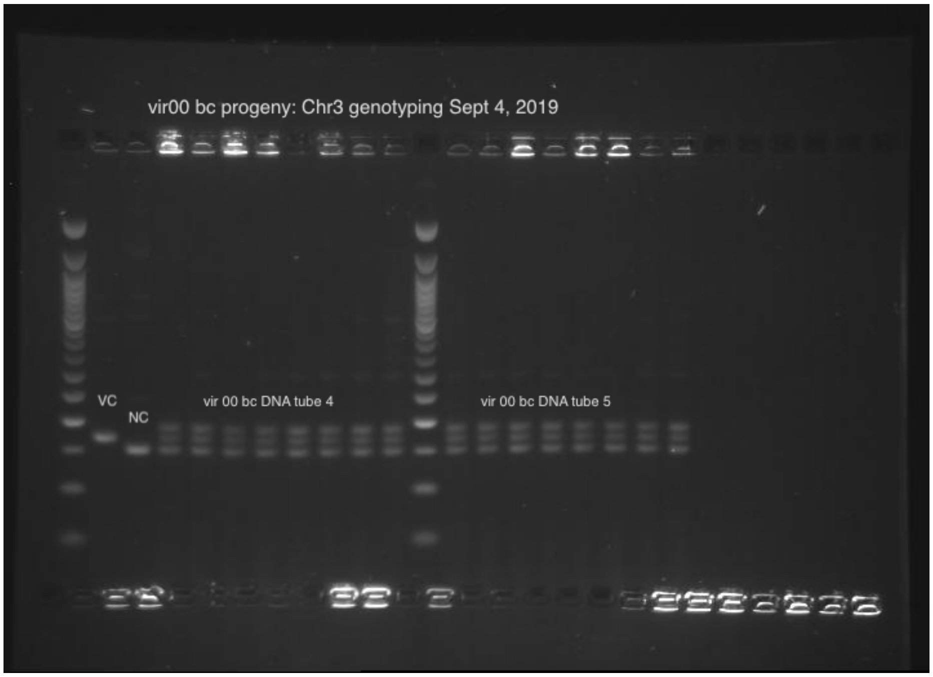


**Figure S2**. Sample genotyping results for Chr3. The first two lanes after the ladder are controls for the *D. virilis* and *novamexicana* allele size, respectively.

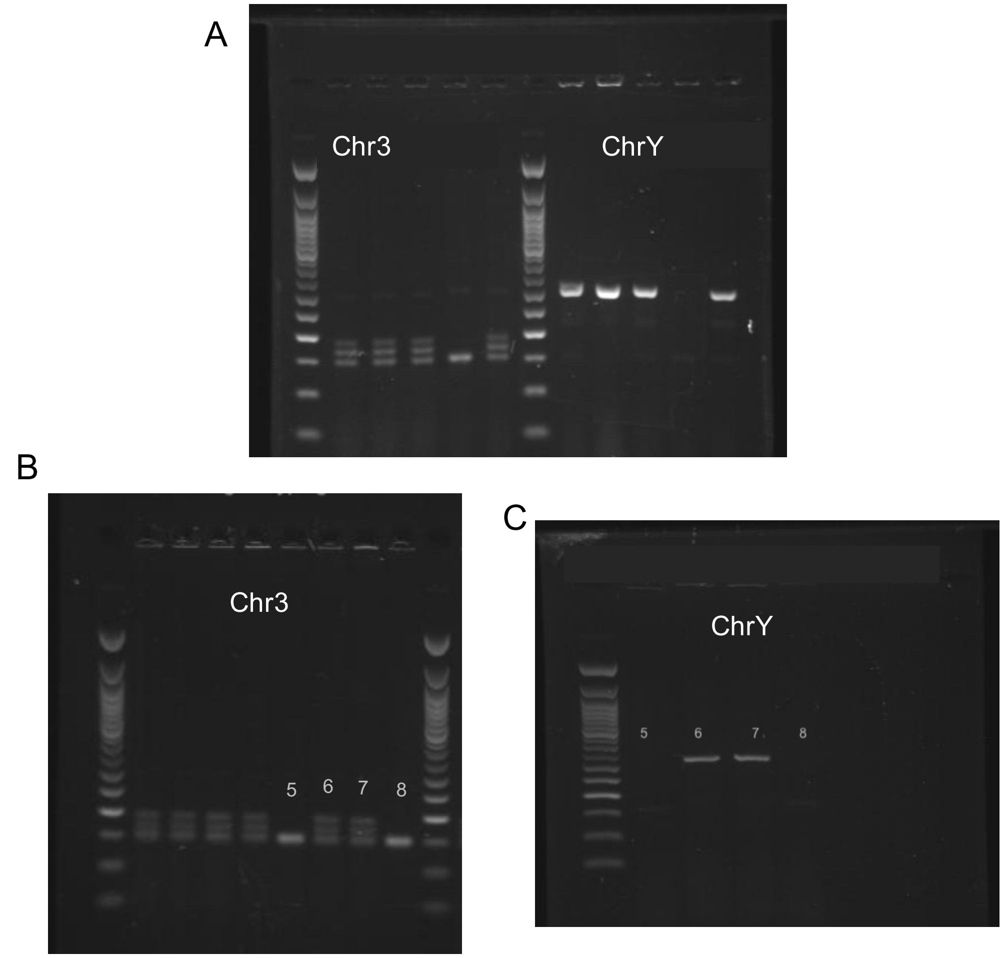


**Figure S3**. Nondisjunction occurred in lines that only had the D.nov Chr3 allele. A) side by side PCR of 5 samples for the Chr3 microsatellite locus (left panel) and ChrY (right panel). Sample 4 has only the D.nov allele and has no Y chromosome, indicating a nondisjunction event. B) PCR of 8 samples for the Chr3 microsatellite locus. Samples 5 and 8 have only the D.nov Chr3 allele. C) PCR of samples 5-8 for ChrY. Samples 5 and 8 have no Y chromosome, indicating a nondisjunction event.

A

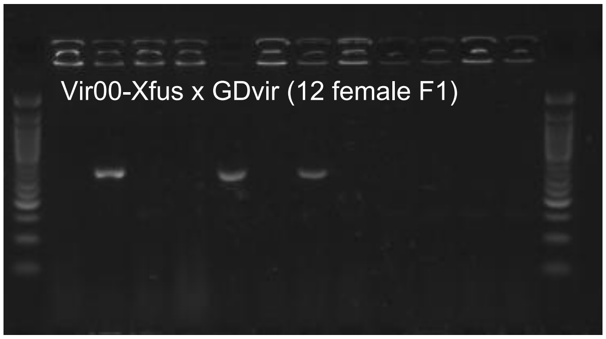


B

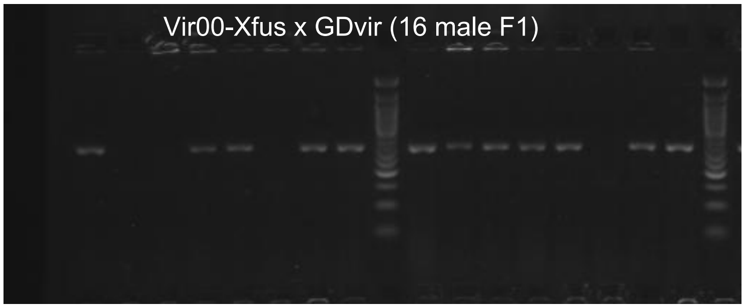


**Figure S5**. X-Y nondisjunction in a vir00-Xfus male. The focal male was crossed to two virgin Gdvir females. A) 3/12 female progeny contain a Y chromosome (XXY karyotype), indicating XY sperm from the father. B) 4/16 male progeny do not contain a Y chromosome (XO karyotype), indicating nullisomic sperm from the father.
