## Supplementary Tables for "Three recent sex chromosome-to-autosome fusions in a *Drosophila virilis* strain with high satellite content"

**Table S1.** Summary of vir00 substrains used for different experiments.

| Substrain | Indel validation | Genetic determination of fusion partner | Resequencing | Nondisjunction assay | DNA damage assay |
| --- | --- | --- | --- | --- | --- |
| Vir00-Yfus | yes | Yes; Chr3 | Yes | Yes | Yes |
| Vir00-Xfus-a | yes | Yes; Chr4 | Yes | Yes | No |
| Vir00-Nofus/Xfus-a | yes | N/A | Yes | No | Yes |
| Vir00-Nofus | yes | N/A | No | Yes | No |
| Vir00-Yfus+Xfus-b | yes | No, unknown autosome is fused to X. | No | No | No |

**Table S2**. Summary of crosses for the Y fusion validation experiment. *D. novamexicana* females were crossed to vir00-Yfus males, and the male F1 progeny were backcrossed to *D. novamexicana* females. Each F2 male progeny were genotyped at each of the 4 candidate autosomal markers which are polymorphic between virilis and novamexicana. The null hypothesis was that 50% male F2s should be homozygous and 50% should be heterozygous. The chi-square p-value was only significant for vir00-Yfus on the Chr3 marker. We included crosses with vir08 instead of vir00-Yfus as a negative control. P-value indicates the p-value for a chi-square test with 1 degree of freedom. The three individuals found to be homozygous for the Chr3 marker actually had a nondisjunction event and did not contain the Y chromosome (Figure S3).

| **Strain** | **Chr2 het--hom** | **Chr3 het--hom** | **Chr4 het--hom** | **Chr5 het--hom** | **p-value (Chr3)** |
| --- | --- | --- | --- | --- | --- |
| vir00-Yfus | 10 -- 8 | 79 -- 3 | 9 -- 9 | 8 -- 10 | < .00001 |
| vir08 (control) | 13 -- 8 | 9 -- 13 | 8 --14 | 14 -- 8 | 0.39 |

**Table S3**. Summary of crosses for the X fusion validation experiment. Vir00-Xfus females were crossed to each of three GFP-insertion lines (vir121, vir117, vir95). The resulting F1 males were crossed to GDvir (virilis genome strain vir87) females. Female progeny were scored for presence or absence of GFP signal in the larval brain. The null hypothesis was that 50% female F2s should be GFP+ and 50% should be GFP-. We included a cross with GDvir instead of vir00-Xfus as a negative control. Chrom indicates the chromosome on which the GFP insertion is located, mapped in Stern et al. 2017. P-value indicates the p-value for a chi-square test with 1 degree of freedom.

| **Cross** | **Chrom** | **GFP + females** | **GFP- females** | **p-value** |
| --- | --- | --- | --- | --- |
| vir00-Xfus x vir121 | 2 | 53 | 70 | 0.125 |
| vir00-Xfus x vir117 | 5 | 103 | 97 | 0.67 |
| vir00-Xfus x vir95 | 4 | 25 | 279 | < 0.00001 |
| Gdvir x vir95 | 4 | 56 | 55 | 0.92 |

**Table S4**. Strains used in this study. Strains were obtained from the Cornell Drosophila Species Stock Center.

| **Strain number** | **Shortform** | **Species** | **Origin** | **Data** |
| --- | --- | --- | --- | --- |
| 15010-1051.51 | vir51 | *D. virilis* | Chile | sequencing |
| 15010-1051.52 | vir52 | *D. virilis* | Russia | sequencing |
| 15010-1051.86 | vir86 | *D. virilis* | Mexico | sequencing |
| 15010-1051.47 | vir47 | *D. virilis* | China | sequencing |
| 15010-1051.49 | vir49 | *D. virilis* | Argentina | sequencing |
| 15010-1051.85 | vir85 | *D. virilis* | Japan | sequencing, comet assay |
| 15010-1051.08 | vir08 | *D. virilis* | California, USA | sequencing, comet assay |
| 15010-1051.00 | vir00 | *D. virilis* | California, USA | sequencing, comet assay, chromosome fusions |
| 15010-1051.118 | vir118 | *D. virilis* | Rwanda, Africa | sequencing |
| 15010-1051.48 | vir48 | *D. virilis* | Mexico | sequencing, comet assay |
| 15010-1051.87 | vir87/GDvir | *D. virilis* | Japan | sequencing, comet assay |
| 15010-1051.09 | vir9 | *D. virilis* | Japan | sequencing, comet assay |
| 15010-1051.121 | vir121 | *D. virilis* | Unknown | X-fusion crossing validation; genotype: Dvir\y[40a]ec[1]cv[1]v[1]si[2]dy[1]w[1]ap[40e];PBac{GreenEye.UAScnnEGFP}Dvir2 |
| 15010-1051.117 | vir117 | *D. virilis* | Unknown | X-fusion crossing validation; genotype: Dvir\y[40a]ec[1]cv[1]v[1]si[2]dy[1]w[1]ap[40e];PBac{GreenEye.UAStubEGFP}Dvir10 |
| 15010-1051.95 | vir95 | *D. virilis* | Unknown | X-fusion crossing validation; genotype: Dvir\y[40a]ec[1]cv[1]v[1]si[2]dy[1]w[1]ap[40e];PBac{5PBlueEye}Dvir1 |
| 15010-1031.14 | nov14/Gdnov | *D. novamexicana* | Moab, Utah. | Y-fusion crossing validation |

**Table S5**. Primer sequences used in this study.

| **Primer name** | **Forward_seq** | **Reverse_seq** | **Purpose** |
| --- | --- | --- | --- |
| chr2_2 | TGGAAATTTCGAGTGGTTCG | GCAAACAGTCAAGCTCGTTTC | To amplify a microsatellite locus that is polymorphic between *D. virilis* and *D. novamexicana* (Y fusion genetic validation) |
| chr3_2 | GTGCAGCAGCCAACAGTC | ATGTCACTCACATGGCCAAA | To amplify a microsatellite locus that is polymorphic between *D. virilis* and *D. novamexicana* (Y fusion genetic validation) |
| chr4_2 | ATTGTGCGAGTCCAGAGTCC | CATTTTGGGAGATGGCAGAC | To amplify a microsatellite locus that is polymorphic between *D. virilis* and *D. novamexicana* (Y fusion genetic validation) |
| chr5_1 | ACACACACGACCCACCACT | GCAGCCAAGTGTTTGCTATG | To amplify a microsatellite locus that is polymorphic between *D. virilis* and *D. novamexicana* (Y fusion genetic validation) |
| vir_Y | CGTCATCCGTTTTGCCAGTG | AGGCATCCCGTATTAAGCGG | Y chromosome genotyping for presence/absence, designed by Yasir Ahmed-Braimah |
| Chr5_indel | CGCATGAACGGACAGGGTAT | AGTCAGTCTTTCTAGCAGGCG | indel genotyping to validate vir00 sublines |
| Chr6_indel | TGTATTCAATTTCGCGACCCT | TCCGCTTTGAGATTGGCCAA | indel genotyping to validate vir00 sublines |
| Chr3_indel | GCGTTATGATGCCAGGACTG | AGCAAAATGGTGGCACTAACT | indel genotyping to validate vir00 sublines |
| Chr2_indel | TGGACTATGCGTGTGAAGGA | CACGGACTTAAATACCTCCTTGA | indel genotyping to validate vir00 sublines |

**Table S6**. Satellite DNA abundance in copy number in the vir00 substrains.

| **Substrain** | **AAACTAC** | **AAATTAC** | **AAACTAT** |
| --- | --- | --- | --- |
| **Vir00-Yfus (rep 1)** | 5645000 | 1648106 | 2207984 |
| **Vir00-Yfus (rep 2)** | 5376774 | 1624806 | 2438514 |
| **Vir00-Xfus** | 5502968 | 1508402 | 2342352 |
| **Vir00-Nofus** | 4881613 | 1500037 | 2041711 |

**Table S7**. Olive moment measurements in all cells used for statistical analysis in Figure 3A. This table is large so we included as a separate csv file.

**Table S8**. Estimated rDNA copy number from read mapping of vir00 substrains and a control strain, vir08.

| Substrain | Estimated rDNA copy number |
| --- | --- |
| Vir00-Yfus (rep 1) | 625.2 |
| Vir00-Yfus (rep 2) | 634.7 |
| Vir00-Xfus | 577.8 |
| Vir00-Nofus | 637.0 |
| Vir08 | 550.4 |
